# Genomic analysis reveals geography rather than culture as the predominant factor shaping genetic variation in northern Kenyan human populations

**DOI:** 10.1101/2021.10.29.466304

**Authors:** Angela M. Taravella Oill, Carla Handley, Emma K. Howell, Anne C. Stone, Sarah Mathew, Melissa A. Wilson

## Abstract

**Objectives:** The aim of this study was to characterize the genetic relationships within and among four neighboring populations in northern Kenya in light of cultural relationships to understand the extent to which geography and culture shape patterns of genetic variation.

**Materials and Methods:** We collected DNA and demographic information pertaining to aspects of social identity and heritage from 572 individuals across the Turkana, Samburu, Waso Borana, and Rendille of northern Kenya. We sampled individuals across a total of nine clans from these four groups and, additionally, three territorial sections within the Turkana and successfully genotyped 376 individuals.

**Results:** Here we report that geography predominately shapes genetic variation within and among human groups in northern Kenya. We observed a clinal pattern of genetic variation that mirrors the overall geographic distribution of the individuals we sampled. We also found relatively higher rates of intermarriage between the Rendille and Samburu and evidence of gene flow between them that reflect these higher rates of intermarriage. Among the Turkana, we observed strong recent genetic substructuring based on territorial section affiliation. Within ethnolinguistic groups, we found that Y chromosome haplotypes do not consistently cluster by natal clan affiliation. Finally, we found that sampled populations that are geographically closer have lower genetic differentiation, and that cultural similarity does not predict genetic similarity as a whole across these northern Kenyan populations.

**Discussion:** Overall, the results from this study highlight the importance of geography, even on a local geographic scale, in shaping observed patterns of genetic variation in human populations.

## Introduction

Among human populations, both geography and culture contribute to modifying patterns of genetic variation. Gene flow can be constrained by geographic distance (Wright 1943). In humans, it is commonly observed that as geographic distance between populations increases, genetic similarity decreases (e.g.,(Manica, Prugnolle, & Balloux, 2005; Novembre et al., 2008; Ramachandran et al., 2005). In addition to geography, genetic variation and population structuring are also influenced by cultural factors, like language (e.g., (Hunley et al., 2008; Nettle & Harriss, 2003; Pagani et al., 2012; Sun et al., 2013; Xu et al., 2010)) or social organization (Bose, Platt, Parida, Drineas, & Paschou, 2021; Chaix et al., 2007; Heyer et al., 2009; Marchi et al., 2017). Like geographic distance, linguistic distance has been shown to correlate with genetic distance (Cavalli-Sforza, Piazza, Menozzi, & Mountain, 1988; Nettle & Harriss, 2003).

Large scale research efforts have aimed to curate genetic variation across some African populations to understand population history and human health and disease (e.g., (1000 Genomes Project Consortium et al., 2015; Choudhury et al., 2020; Gurdasani et al., 2015; Mulindwa et al., 2020; Tishkoff et al., 2009)). However, with more than 2,000 ethnolinguistic groups across the continent, there is still much to learn about the determinants of substructure both among and within individual ethnolinguistic groups at smaller geographic scales. For example, do cultural boundaries play an important role in shaping gene flow among neighboring groups, or does geography primarily shape patterns of genetic variation, even on a local scaleã

The Turkana, Samburu, Waso Borana, and Rendille are neighboring pastoral ethnolinguistic groups inhabiting the semi-arid northern region of Kenya. They herd cattle, camel, sheep and goat, and migrate over varied distances to access pasture and water. There is intense competition for scarce dry season grazing and water resources in this region, and armed livestock raiding occurs especially between communities belonging to different ethnolinguistic groups (Handley & Mathew, 2020). Marriage across ethnic boundaries particularly between the Rendille and Samburu has been noted (Spencer, 2012). Nomadic pastoralism in all four populations involves lots of movement of people over large distances, including into one another’s territory. While there are no cultural prescriptions against intermarriage between these groups, historical and contemporary livestock raiding and resource competition often lead to hostile relationships between these groups. It is currently unclear whether these tensions contribute to isolation among these groups, or whether gene flow occurs despite the socio-political barriers.

All four groups have lineage-based divisions where individuals are organized into clans and, for some groups, clans are further grouped into either moieties or phratries (**Figure S1**). The Turkana, Samburu and Rendille are exogamous at the clan level while the Borana are exogamous at the moiety level, so for these groups, individuals generally marry outside of their birth clan. In addition to lineage-based divisions, the Turkana are unique from the other three groups in that they also have territory-based divisions that cross-cut clan-level organization. There are no marriage restrictions at the territorial level. This territorial division provides an opportunity to investigate whether territory-based division impacts substructuring in the Turkana. Detailed descriptions of the social organization of these populations can be found in (Handley & Mathew, 2020).

Each of these groups has a patrilineal descent system where an individual’s natal clan affiliation typically follows that of their father. There are, however, situations in which children do not take on the clan identity of their biological father. When the biological father is not officially married, i.e., has yet to pay a bride price to the family of the child’s mother at the time of birth, the child remains affiliated with the natal clan, which is the child’s mother’s and maternal grandfather’s clan. Households with few children or that have relatively higher material security may adopt a child, in which case the child takes the clan identity of the adoptive father. Therefore it is unclear how cultural systems of patrilineal descent shape patterns of male-specific genetic variation within these groups.

In a previous study, two members of this research team sampled 750 individuals from nine clans across these four ethnic groups, and in the Turkana additionally included three territorial sections, to obtain data on cultural beliefs and norms, and to quantify levels of cultural differentiation (cultural F_ST_) among these groups (Handley & Mathew, 2020). Cultural F_ST_ is the proportion of the total variation in cultural traits that lie between populations (Bell, Richerson, & McElreath, 2009; Handley & Mathew, 2020) and can provide a quantitative measure of cultural similarity between groups. For the current study, we sampled individuals from these same nine clans and three Turkana territorial sections, which allows us to examine the relationship between genetic and cultural differentiation.

To form a better understanding of the population structure among the Turkana, Samburu, Rendille, and Waso Borana ethnolinguistic groups and how geography and culture contribute to shaping genetic variation in northern Kenya, we worked with these local groups to obtain genetic samples from 572 individuals across all four populations (**Table 1**). We were able to successfully genotype 376 of the 572 individuals on Illumina’s Multi-Ethnic Global Array (**Table S1**). For all samples, we additionally collected culturally relevant demographic information that included natal and post-marital affiliations and spoken languages for themselves, parents, and grandparents. For married men, we additionally collected demographic information for their spouse(s) (e.g. spouse’s ethnic group and natal clan affiliation, etc.). We report here that geography predominately shapes genetic variation within and among human groups in northern Kenya. Specifically, we found a clinal pattern of genetic variation that mirrors the overall geographic distribution of the individuals we sampled. We found evidence of gene flow and relatively higher rates of intermarriage between the Samburu and Rendille than between any other pair of groups in our sample. We further observed strong recent genetic substructuring among the Turkana, based on territorial section affiliation, that did not affect the between-ethnolinguistic group comparisons. Within ethnolinguistic groups, we found that male Y chromosome haplotypes do not consistently cluster by natal clan affiliation. Finally, we found that ethnolinguistic groups that are geographically closer have lower genetic differentiation, and that cultural similarity (estimated via cultural F_ST_) does not predict genetic similarity as a whole across these four northern Kenyan populations. Overall, despite cultural and linguistic differences, our analysis suggests that geography is the main driving force of genetic variation, even on a very local geographic scale.

**Table 1.** Background information on study populations and collected samples. We sampled a total of 572 individuals in northern Kenya. Here, we describe general background information on the four study populations, including information on language, social organization, and population sizes in Kenya. This table was adapted from (Handley and Mathew 2020) but altered to reflect sample sizes collected in the current study. ^a^Language information reported here is based on assignments from the *World Atlas of Languages (WALS) Online*. ^b^Population sizes reported here were obtained from the 2019 census report of the *Kenya National Bureau of Statistics*. ^c^Borana extend into Ethiopia so their total population exceeds the numbers living in Kenya. ^d^Sample numbers are based on natal affiliations. Since we opportunistically sampled in these regions, we also sequenced individuals beyond the targeted clans and territories and these samples are marked as “Other” here. One individual was from the Gabbra ethnic group and not included in this table.

| Ethnolinguistic group | Language genus<sup>a</sup> | Spoken language | Social organization | Approx. population in Kenya<sup>b</sup> | Clans sampled (sample size)<sup>d</sup> |
| --- | --- | --- | --- | --- | --- |
| Borana | Lowland East Cushitic | Southern Oromo | 2 exogamous moieties<br>17 clans | 276,236<sup>c</sup> | Noonituu (40)<br>Warrajidaa (40)<br>Other (30) |
| Rendille | Lowland East Cushitic | Kirendille | 2 phratries<br>9 exogamous clans | 96,313 | Ldupsai (43)<br>Saale (45)<br>Other (22) |
| Samburu | Nilotic | Northern Maa | 2 phratries<br>8 exogamous clans | 333,471 | Lpisikishu (40)<br>Lukumai (42)<br>Other (27) |
| Turkana | Nilotic | Kiturkana | 18 territorial sections (TS)<br>24 exogamous clans cross-cutting the TS | 1,016,174 | Kwatela TS:<br>Ngisiger (24),<br>Ngipongaa (21),<br>Ngidoca (23)<br>Ngiyapakuno TS:<br>Ngisiger (24),<br>Ngipongaa (26),<br>Ngidoca (22)<br>Ngibochoros TS:<br>Ngisiger (21),<br>Ngipongaa (22),<br>Ngidoca (21)<br>Other (38) |

## Materials and Methods

### Community engagement and ethics

Both SM and CH have worked in Northern Kenya for over a decade and have established and maintained a strong relationship with the local communities. Research with the Turkana, Borana, Rendille, and Samburu has expanded to include genetic analyses and great care has been taken to ensure ethical informed consent, data collection, outreach communication, and data sharing.

Subsequent to obtaining the appropriate research permitting through Kenya’s National Commission for Science, Technology and Innovation (NACOSTI), yet prior to commencing any data collection, the field teams spent a considerable period of time with each local community and its leaders to sensitize individuals to the purpose and process of collecting genetic samples for this study. Other than SM and CH, field teams were composed solely of individuals from the participant communities, many of whom had been working with SM and CH for several years. Research assistants (RAs) and guides worked within their own ethnic groups, therefore having one team per group, and all information was presented to participants using local languages. As a key goal of any informed consent process, potential subjects must demonstrate sufficient comprehension of the methods and underlying scientific principles on which to base their decision to participate. With literacy rates for northeastern Kenya estimated below 10% of the adult population (Kebathi, 2008) explaining fundamental concepts regarding DNA, genetic data sample collection, and data sharing was of paramount importance. For each area surveyed, teams met with the responsible county/deputy commissioners, local/assistant chiefs, and/or community elders councils to explain the purpose of the study and to obtain permissions from the appropriate bodies. As data collection began, RAs discussed the study, its purpose, methodologies, and underlying scientific principles to each eligible participant and provided ample time for participants to ask questions and address any concerns. At this time, there was an estimated drop-out rate of 20% of eligible participants. For those remaining, we transitioned to the formal consent process, where the objectives, methods, and benefits of the study would be repeated before asking a subject to sign or mark the consent form. At this stage, there was an additional estimated 15% drop-out of eligible participants. Those who agreed to take part in the study were provided with the contact information for both local and foreign research team members in the event that s/he should choose to be removed from the study at any future point. Furthermore, despite the common practice in many locations of husbands granting permission for themselves and for their wives, permissions were obtained explicitly and directly from all female participants. However, the research teams made efforts to avoid households where directly soliciting female participation could transgress cultural norms and inadvertently introduce additional domestic concerns.

Once obtaining consent for participation, research assistants demonstrated the cheek cell swab collection procedure, using a clean swab on themselves to scrub the inside of their own cheeks. Our initial intention was for RAs to swab the participants’ mouths; however, we found that participants felt more comfortable being in control of the process to swab their own mouths. This required oversight from the RAs to ensure that the swab was oriented in the appropriate direction, did not come into contact with foreign bodies, and that enough pressure and effort were applied during the collection. Furthermore, we requested that participants rinse their mouths with water if they recently had been chewing tobacco or other organic products. Satisfactory swabs were handed to the RAs to seal within the collection tube and returned to CH for cool bag storage. After sample collection, participants were asked to respond to a 10-15 minute survey, developed in ONA and implemented in the field through Online Data Kit (ODK) using handheld Samsung tablets. The survey requested permission to record the GPS locations of participants along with questions regarding the biological and cultural kinship lineages of the participants with a resolution to both maternal and paternal grandparents, along with languages spoken within each family household.

In 2018, AMTO traveled to Turkana to present outreach materials and discuss initial findings. We worked with VizLab graphic design studio at ASU to make a series of images to help explain what DNA is, how to get DNA from a cell, what can be learned about people and human history with DNA, and preliminary results from this genetic study (**Figure S2**). Along with the images, we created a script to explain, in layman’s terms, what the images mean; we also had questions to ask at the end of the presentation to make sure participants were following and understanding what we were demonstrating to them. We worked with the local field assistants to translate the script into the local language and to present the script to the community. We presented to a total of 6 settlement areas across 3 territorial sections in Turkana County, Kenya. Overall, the presentations were well received by the communities, and people expressed interest in the results. Some people also expressed their excitement about wanting to know what else we would find from their DNA. Additional dissemination from our group will occur in the near future as it becomes safer to travel.

The Turkana, Rendille, Samburu, and Borana are small-scale pastoral populations in northern Kenya. We are therefore taking measures to ensure the protection of these groups by providing the genetic data generated here as controlled access while maintaining appropriate standards of data access. The genomic data generated here will be available through dbGap.

### Sampling and sequencing

We collected DNA and demographic information from a total of 572 individuals from Turkana, Samburu, Rendille, and Waso Borana and successfully genotyped 376 of these individuals on Illumina’s Multi-Ethnic Global Array (**Table 1; Table S1**). For each ethnolinguistic group, we sampled individuals from at least two clans. Data collection occurred across northern Kenya from October 2016 – October 2017. For each participant, a DNA sample was taken in the form of saliva or cheek swab; the saliva was collected in an Oragene OG-500 DNA collection kit. In addition to collecting a DNA sample, a questionnaire was administered to each participant to acquire demographic information; this information included, for example, natal and post-marital clan affiliation, and spoken languages. DNA was extracted using a phenol-chloroform extraction method for the samples collected from cheek swabs. DNA for the samples collected with the Oragene OG-500 DNA collection kit was extracted at Yale Center for Genomic Analysis. The extracted DNA was then quantified on both a Qubit and Nanodrop. Each sample’s extracted DNA was then diluted to at least 35 ng/ul in a volume of 40 ul and sent to Langebio-Cinvestav sequencing facility in Mexico for SNP genotyping on Illumina’s Multi-Ethnic Global Array.

### Quality control and filtering

We received the SNP genotype data in the form of a raw plink file. The coordinates were mapped to the human reference genome hg19. Initially, there were a total of 1,779,819 markers genotyped on the array. Sites with no valid mapping for the probe or with more than 1 best-scoring mapping for the probe were removed from our analyses. Additionally, we removed any sites marked as insertions or deletions. There were 27,089 duplicated variants in this file; duplicated variants have the same chromosome number and position and can have the same or different allele codes. Duplicated variants with the same chromosome, position, and allele codes were merged, while duplicated variants with the same chromosome and positions, but different allele codes were removed. We merged the duplicated sites using the ‘--merge-equal-pos’ flag and the default merge mode - which ignores missing calls and sets mismatching genotypes to missing - using PLINK v1.9 (Chang et al., 2015). There were a total of 1,715,718 sites after filtering.

As an additional quality control measure, we inferred the sex chromosome complement of each individual and compared this information with reported sex information. Two approaches were used to infer the sex chromosome complement of each individual, one approach based on the X chromosome inbreeding coefficient (F) and the other approach based on the number of Y chromosome genotype counts. Since genetic males are expected to have one X chromosome, they should not have any heterozygous sites on the X chromosome (minus the pseudoautosomal regions - PARs) and therefore an inbreeding coefficient equal to 1. We used the “--check-sex ycount” flag in PLINK v1.9 (Chang et al., 2015) to calculate the X chromosome inbreeding coefficient and the number of Y chromosome genotype counts using. The PARs were excluded from this calculation.

As per the PLINK documentation recommendations, individuals with an X chromosome inbreeding coefficient greater than 0.8 were considered male, while individuals with an X chromosome inbreeding coefficient less than 0.2 were considered female. The expectation for genetic females is that they have no genotype calls on the Y chromosome - since genetic females are expected to be XX - however, all the female individuals in our data set had genotype calls on the Y chromosome. We, therefore, visualized the X chromosome inbreeding coefficient and non-missing Y chromosome genotype counts together to see the distribution of these values in males and females and to identify any individuals that did not cluster with expected male and female values. We removed individuals that had discrepancies between these two metrics (**Figure S3**). A total of 10 individuals were removed.

Identity by descent (IBD) was calculated across the autosomes to identify and remove related individuals. Prior to running the IBD analysis, we filtered sites with missing data across samples greater than 5% (--geno 0.05 flag in PLINK), sites with Hardy-Weinberg equilibrium p-value threshold less than 1×10^−50^, and pruned sites for linkage disequilibrium (50 kb window size, 10 kb variant step size, 0.2 r^2^ threshold). Filtering and IBD were calculated, using PLINK v1.9 (Chang et al., 2015), for all pairwise combinations of samples in our data set and output pairs of individuals with more than 18% IBD. We removed two samples with 100% IBD that were not replicate samples (the same sample sequenced twice), as these may reflect contamination. We had two replicates in this data set and removed one sample from each replicated pair. In the cases where there were clusters of individuals related, we removed the individuals who were related to many other individuals, to minimize the number of individuals to remove. In cases where just two individuals were related, we attempted to remove roughly equal numbers of males and females when possible. A total of 67 samples were removed, leaving 301 individuals.

We next performed an initial principal components analysis (PCA) on the 301 samples using smartpca, a program within the EIGENSOFT v6.0.1 software package (Price et al., 2006). We identified a total of 4 outlier samples; these samples were removed from subsequent analyses (**Figure S4**).

After individual filtering, we performed site filtering on the autosomes, Y chromosome, and mitochondrial DNA. For the autosomes, we removed sites with more than 5% missing data across individuals at a given site (“--geno 0.05” flag in PLINK/ 95% call rate filter), removed sites that deviate from Hardy-Weinberg equilibrium (p-value threshold of 1×10^−50^), and performed linkage disequilibrium pruning (50 kb window size, 10 kb variant step size, and 0.2 r^2^ threshold). For the Y chromosome and mitochondrial DNA, we removed sites with heterozygous calls, removed sites with more than 5% missing data across individuals at a given site, and removed sites that deviate from Hardy-Weinberg equilibrium with a p-value threshold of 1×10^−50^. After filtering, 516,821, 3,295, and 811 sites remained on the autosomes, Y chromosome, and mitochondrial DNA, respectively.

### Population genetic analyses

To explore the genetic structure within and among northern Kenyan populations, we ran PCA using smartpca (Price et al. 2006) and ADMIXTURE (Alexander, Novembre, & Lange, 2009) on the autosomes for all unrelated samples. We ran ADMIXTURE for K = 2 - 5 with a total of 10 replicates for each K value. For streamlined post analysis and visualization of the different replication runs and K values from the ADMIXTURE analysis, we used pong (Behr, Liu, Liu-Fang, Nakka, & Ramachandran, 2016), an algorithm for processing and visualizing membership coefficient matrices. Pong finds the best alignment across all runs within and across the different K values and identifies modes among all runs for each K. We used the best alignments across all runs within and across the different K values for visualization in this manuscript.

To quantify genetic differentiation within and among northern Kenyan populations, we calculated Hudson’s *F*_ST_ (Hudson, Slatkin, & Maddison, 1992) using the estimator derived in (Bhatia, Patterson, Sankararaman, & Price, 2013). *F*_ST_ was calculated on the autosomes for ethnolinguistic groups and Turkana territorial sections. Results were visualized in R (R. C. Team & Others, 2013; R. Team & Others, 2015), using the visualization package, ggplot2 (Wickham, 2011).

To test whether genetic *F*_ST_ was different between Ngibochoros and at least one of the ethnolinguistic groups and Kwatela and/or Ngiyapakuno and the same ethnolinguistic group, we performed a series of permutations. We randomly shuffled samples from two of the territorial sections, calculated *F*_ST_ between the territorial section and ethnolinguistic group, and then calculated the test statistic which was the absolute difference between *F*_ST_ for each territorial section. We repeated this 1,000 times and calculated the p-value. This was done for all combinations.

To visualize the relationships among haplotypes within each ethnolinguistic group, we generated haplotype networks for the Y chromosome and mitochondrial DNA. SNPs in our data set were first set to the forward orientation. Sites on the opposite strand were identified using snpflip (https://github.com/biocore-ntnu/snpflip) and flipped using PLINK v1.9 (Chang et al., 2015). Snpflip uses information from a reference genome fasta sequence and the bim PLINK file to identify SNPs on the reverse strand. The GRCh37 GENCODE reference genome was used (ftp://ftp.ebi.ac.uk/pub/databases/gencode/Gencode_human/release_30/GRCh37_mapping/GRCh37.primary_assembly.genome.fa.gz). Any sites unable to be flipped due to ambiguity in whether the site was on the reverse strand were removed. A total of 2,807 and 806 sites remained on the Y chromosome and mitochondrial DNA, respectively. PLINK files were then converted to VCF format using PLINK v.1.9 (Chang et al. 2015). VCF files were converted to FASTA file format using python and haplotype networks were constructed in R (R. C. Team & Others, 2013; R. Team & Others, 2015) using pegas (Paradis, 2010) and ape (Paradis & Schliep, 2019) packages.

### Quantification of intermarriage

To quantify the amount of intermarriage in the Turkana, Samburu, Rendille, and Waso Borana, we used questionnaire information collected here and from (Handley & Mathew, 2020). The questionnaire information we collected in this study requested spouses’ ethnolinguistic group information only for the married men, while the questionnaire information from (Handley & Mathew, 2020) had spouse information for both married men and women that were sampled. In these groups, men may marry more than one wife, so rates of marriages were based on the total number of marriages rather than the number of individuals. For each ethnolinguistic group, we calculated the percentage of marriages both within the same ethnolinguistic group and from different ethnolinguistic groups. This was calculated separately for men and women.

### Correlations between genetic and cultural differentiation

To test whether measures of genetic similarity can predict cultural similarity, we performed a series of Pearson’s correlations between genetic *F*_ST_, cultural *F*_ST_, geographic distance, and linguistic distance. We used cultural *F*_ST_ values and linguistic distances that were previously calculated among these ethnolinguistic groups (Handley & Mathew, 2020). Briefly, cultural *F*_ST_ is a measure of cultural similarity between two groups; a low cultural *F*_ST_ indicates two groups are more culturally similar while a higher cultural *F*_ST_ indicates two groups are less culturally similar (Bell et al., 2009). For each group, language, genus, and family information were acquired from The World Atlas of Language Structures (WALS) database (“WALS Online - Home,” n.d.). Using this information linguistic distances were categorized as follows: a score of 0 for groups that speak the same language (same language, same genus, same family), a score of 1 for groups that speak different languages from the same language genus (different language, same genus, same family), a score of 2 for groups that speak different languages from different genus but within the same family (different language, different genus, same family), and finally, a score of 3 for groups that speak different languages from different language families (different language, different genus, different family). Further details can be found in (Handley & Mathew, 2020). To calculate geographic distance between groups, we collected GPS coordinate information for the locations in which genetic sampling occurred. If a household fell within one precision of one or more households (within 20 meters), only one GPS measure was recorded. Using the latitude and longitude for each measured household in each population, we calculated the average distance between pairs of populations in kilometers (km). This involved calculating the distance between all households from one population to another population and then averaging these distances. This was computed for all pairs of populations using a custom python script. Pearson correlations were performed in R (R. C. Team & Others, 2013; R. Team & Others, 2015) using package ppcor (Kim, 2015) and visualized using ggplot2 (Wickham, 2011).

### Data and code availability

The genotype data generated in this manuscript has been deposited on dbGap (dbGap accession number phs002654.v1.p1) and will be made available upon publication. All original code used in this manuscript can be found on GitHub: https://github.com/SexChrLab/Kenya_Fst.

## Results

### Intermarriage among ethnolinguistic groups contributes to the clinal pattern of genetic variation

We found a clinal pattern of genetic variation that mirrors the overall geographic distribution of the individuals we sampled (**Figure 1A, 1B**). In the principal components analysis (PCA), the Turkana samples separate from the other three groups along PC1, and along PC2 the Borana samples separate from the Rendille and Samburu (**Figure 1B**). While the Borana samples form a discrete cluster from the other ethnolinguistic groups, there is overlap between Samburu and Rendille and some overlap between Samburu and Turkana (**Figure 1B**). Interestingly, many of the overlapping Samburu and Rendille samples have a family history - a parent and/or grandparent(s) in the Rendille and Samburu, respectively (**Table S2**). In contrast, the Samburu sample that falls near Turkana and the Turkana sample that falls near Samburu have no reported cross-group family history through the grandparent level in these individuals (**Table S2**). We additionally found high rates of intermarriage between some ethnolinguistic groups and nearly non-existent intermarriage between others (**Figure 1C**). For Rendille, 5% of female marriages and 16.4% of male marriages were with a Samburu individual (**Figure 1C**). For Samburu, we observe almost the exact opposite pattern, with 11.4% of female marriages and 4.8% of male marriages with a Rendille individual (**Figure 1C**). The Samburu also have low levels of intermarriage with the Turkana; 3.2% of male Samburu marriages were with a Turkana individual (**Figure 1C**). For the Borana, none of our sampled individuals report marriages with the Turkana, Samburu, or Rendille, but did report varying levels of intermarriage between the Borana and Sakuye, Gabra, Garri, and Somali (**Figure 1C**).

**Figure 1.**
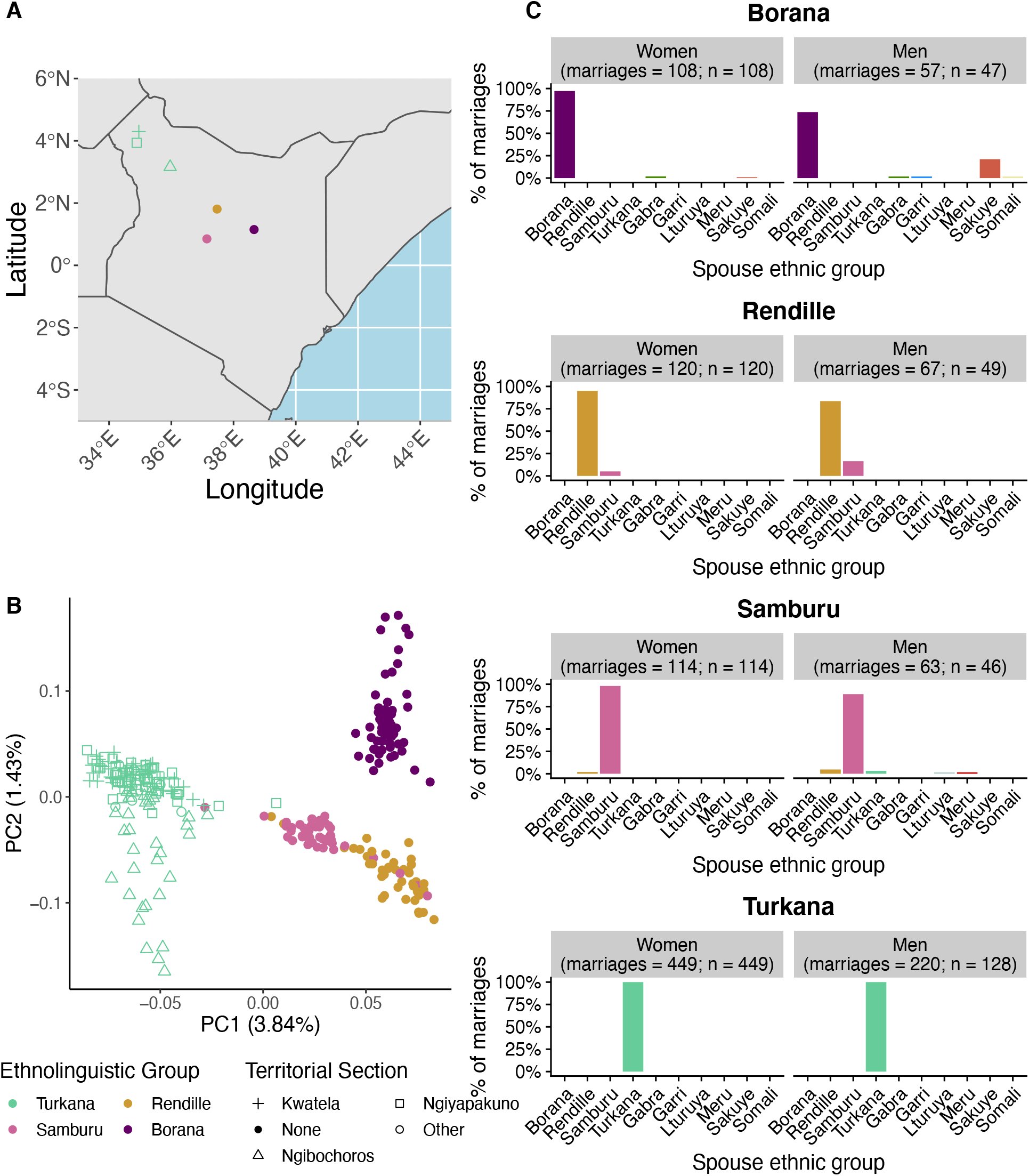
Intermarriage among ethnolinguistic groups contributes to the clinal pattern of genetic variation. Sampling regions, patterns of genetic variation, and rates of intermarriage across northern Kenya human populations. A) We sampled 376 individuals across four ethnolinguistic groups in northern Kenya and for the Turkana only, we additionally sampled across three territorial sections. B) Autosomal principal components analysis (PCA). C) Rate of intermarriage across each ethnolinguistic group. Points in A and B represent sampled ethnolinguistic groups and Turkana territorial sections. Colors represent ethnolinguistic group affiliation, and shapes represent Turkana territorial section affiliation. Each point in A represents the geographic location of each sampled group, while the points in B represent individuals.

### The Turkana have additional variation and geography-based substructuring

Just as these ethnolinguistic groups are geographically separated and similar to the PCA (**Figure 1B**), we observed clear genetic separation in ADMIXTURE analyses (**Figure 2A**). In the ADMIXTURE analyses, each of the ethnolinguistic groups have their own unique ancestry at K = 5 (**Figure 2A**). Interestingly, at K = 4 we observe possible admixture between Samburu and Rendille (**Figure 2A**).

**Figure 2.**
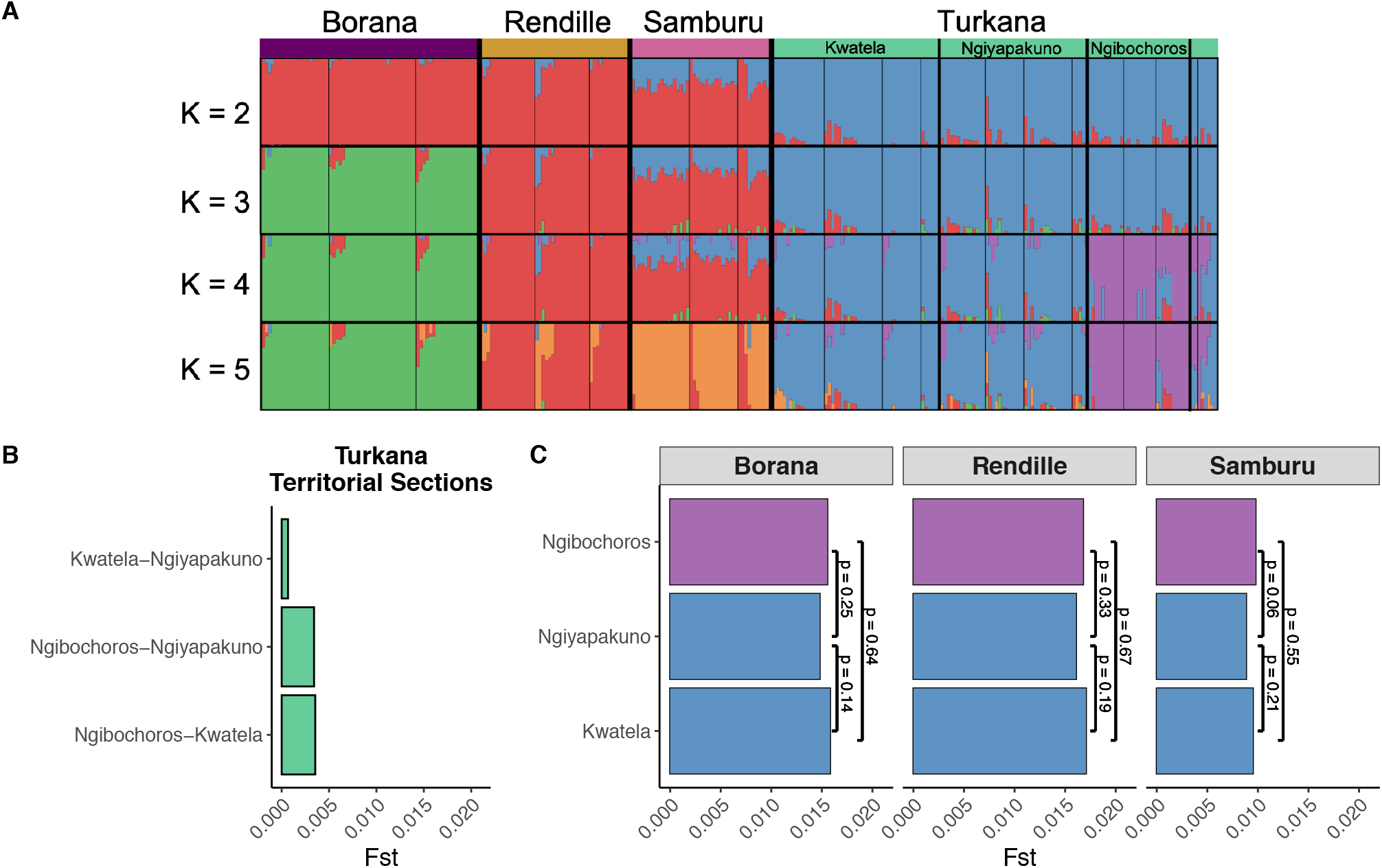
The Turkana have additional variation and geography-based substructuring. A) ADMIXTURE analysis for 10 replicates of K = 2 - 5 for the autosomes. Each vertical bar represents an individual, and the colors represent the proportion of ancestry corresponding to K. Samples are organized by ethnolinguistic groups (separated by thick black vertical bars), then by Turkana territorial sections (separated by medium black vertical bars), and lastly by natal clan affiliation (separated by thin black vertical bars). We observe no substructure based on natal clan affiliation but do observe geographic substructuring in the Turkana based on territorial section (purple and blue clusters at K = 4 and 5). B) Autosomal genetic differentiation (*F*_ST_) among Turkana territorial sections. Individuals from Ngibochoros territorial section are more genetically different than individuals from the other sampled territories. C) Autosomal *F*_ST_ among each Turkana territorial section and the other sampled ethnolinguistic groups. We performed a series of pairwise permutations and found that there is no statistical difference in genetic differentiation among Turkana territorial sections and ethnolinguistic groups. P-values from the permutation tests are annotated on the plot.

In the Turkana, we additionally found geography-based genetic substructuring based on territorial region (**Figure 2A; Figure S5**). In the ADMIXTURE analyses, we see substructure within the Turkana before we see all four ethnolinguistic groups being identified separately. For example, at K = 4, the Turkana are characterized by two different ancestries, with one of these ancestries unique to individuals from the Ngibochoros territorial section (**Figure 2A**). Consistent with this, in the PCA, we observe variation within the Turkana along PC2 (**Figure 1B**). The individuals from the Ngibochoros territorial section separate from the other Turkana territorial sections along PC2. We further calculated genetic differentiation, *F*_ST_, among the three Turkana territorial sections. We found that *F*_ST_ between Ngibochoros and either Kwatela or Ngiyapakuno are much higher than *F*_ST_ between Kwatela and Ngiyapakuno, which are territorial sections that are both adjacent to each other and distant from the Ngibochoros (**Figure 1A; Figure 2B**).

The observed territorial section substructuring in the Turkana may be due to geographic separation among the territorial sections. Alternatively, it is possible that individuals from Ngibochoros - the territorial section that is closer geographically to the Borana, Samburu, and Rendille than the other Turkana territorial sections - may have admixed with the other neighboring groups, resulting in the higher genetic differentiation. To investigate these scenarios further, we calculated genetic *F*_ST_ between each of the Turkana territorial sections and the other three ethnolinguistic groups and performed permutation tests to investigate whether *F*_ST_ values were significantly different between each territorial section and ethnolinguistic group. If gene flow was occurring among Ngibochoros and the other three ethnolinguistic groups, we would expect *F*_ST_ to be lower between Ngibochoros and at least one of the ethnolinguistic groups than between either Kwatela or Ngiyapakuno and the same ethnolinguistic groups. What we find, however, is that *F*_ST_ is not significantly different for each of the Turkana territorial sections when compared with each of the other ethnolinguistic groups (**Figure 2C**).

### Y chromosome haplotypes do not consistently cluster by natal clan affiliation

On the Y chromosome - where we expected Y haplotypes to be more similar for males from the same clan in groups with patrilineal descent than in different clans - we found that haplotypes do not consistently cluster by natal clan affiliation. For Turkana and Borana, there are no haplotypes unique to a clan (**Figure 3A, 3D**). For the Samburu, most of the haplotypes cluster by natal clan affiliation, with the exception of one haplotype that is shared among individuals from both clans we sampled (**Figure 3B**). For the Rendille we observed one haplotype unique to individuals from the Ldupsai clan, however, the rest of the haplotypes were shared among clans (**Figure 3C**). We observe similar characteristics in the mitochondrial DNA haplotype networks (**Figure S6**). Overall, within these patrilineal descent groups, Y chromosome haplotypes generally do not cluster by natal clan affiliation.

**Figure 3.**
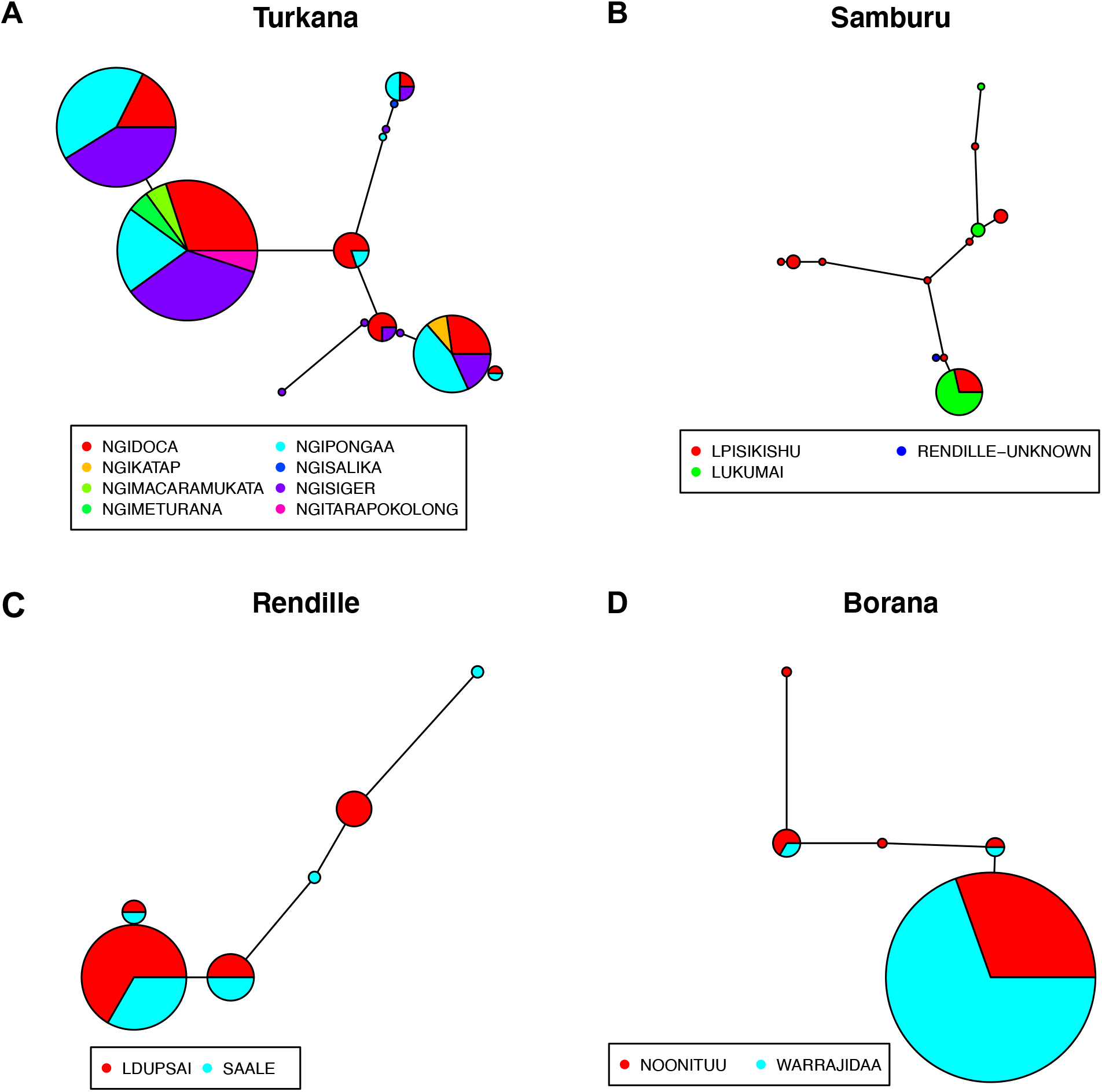
Y chromosome haplotypes do not consistently cluster by natal clan affiliation. Haplotype networks constructed from Y chromosome SNP data from A) Turkana, B) Samburu, C) Rendille and D) Borana male samples. The size of each node (circle) is proportional to the number of samples in the node (larger nodes have more samples and smallest nodes have 1 sample). Colors within each node represent natal clan affiliation corresponding to the key in each panel.

### Genetic differentiation is driven by physical separation, not cultural processes

Ethnolinguistic groups that are geographically closer typically had lower genetic *F*_ST_ (**Figure 4**). The lowest genetic *F*_ST_ was found between the Samburu and Rendille yet they speak languages from different language families; individuals from these ethnolinguistic groups are closer geographically to each other than to any other ethnolinguistic group. The Turkana sampled here are, on average, furthest geographically from the other ethnolinguistic groups - ranging from 286 km to 439 km away. The Turkana also have much higher genetic *F*_ST_ values with the Rendille, Borana, and Samburu than *F*_ST_ measured between any two comparisons of the Rendille, Borana, and Samburu (**Figure 4**).

**Figure 4.**
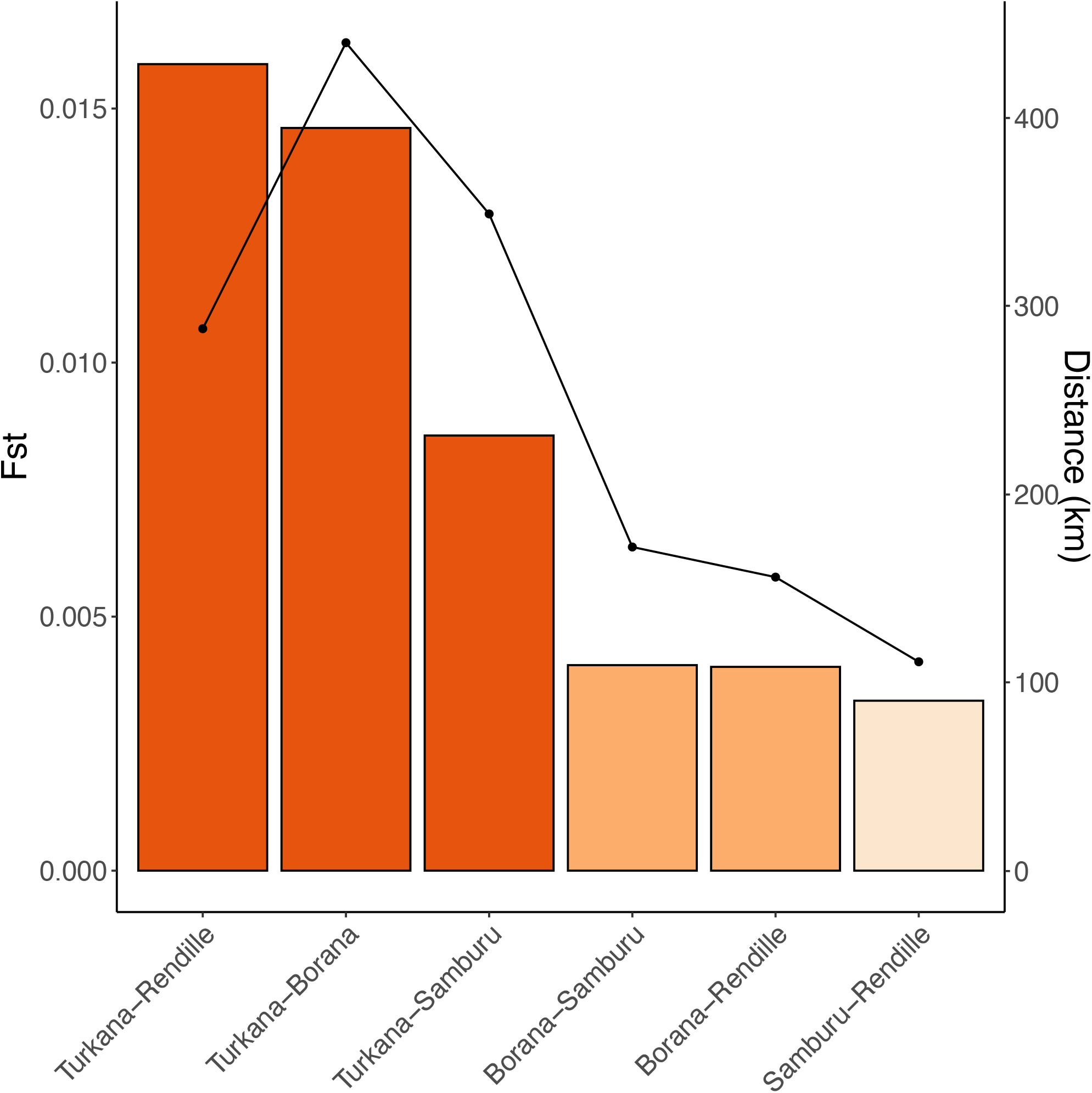
Geography primarily impacts patterns of genetic differentiation among ethnolinguistic groups but for some groups, the pattern of genetic differentiation secondarily parallels linguistic relationships. We calculated autosomal *F*_ST_ among ethnolinguistic groups. Bars are ordered by *F*_ST_. Dark orange (left) are groups furthest geographically, while the lighter orange bars are groups closest geographically. The Samburu and Rendille (pale orange) are two neighboring groups that speak languages from different language families, yet have the lowest genetic *F*_ST_ observed in our study. Line graph corresponds to the geographic distance between each pair of ethnolinguistic groups

For some groups, the pattern of genetic differentiation secondarily paralleled linguistic relationships. Among the Turkana genetic *F*_ST_ comparisons, genetic *F*_ST_ between Turkana and Samburu - both Nilo-Saharan speakers - is about two times lower than genetic *F*_ST_ between the Turkana and Rendille, even though the Rendille individuals sampled here are closer to Turkana by about 63 km (**Figure 4**).

We find that cultural differentiation does not predict genetic differentiation among neighboring groups in northern Kenya. Genetic *F*_ST_ and cultural *F*_ST_ are not significantly correlated with each other (R = 0.63, p-value = 0.18; **Figure 5; Figure S7**). However, we observe a significant positive correlation between genetic *F*_ST_ and geographic distance both at the ethnolinguistic group level and also Turkana territorial section level (ethnolinguistic group level: R = 0.82, p-value = 0.048; Turkana territorial section level: R = 0.992, p-value = <0.001; **Figure 5; Figure S7**).

**Figure 5.**
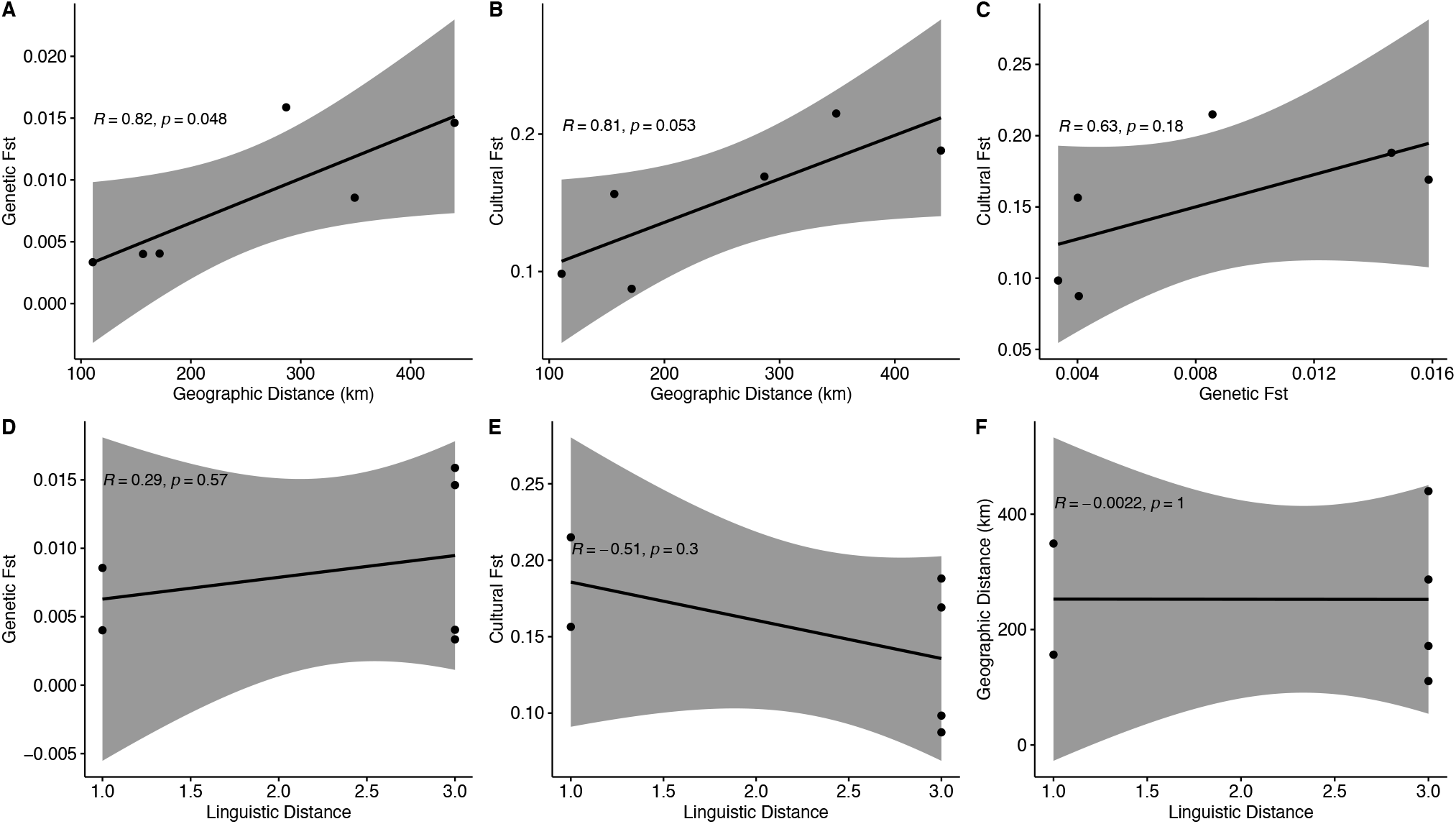
Cultural differentiation does not predict genetic differentiation among human ethnolinguistic groups in northern Kenya. We performed a series of Pearson’s correlations to explore whether cultural differentiation may impact genetic *F*_ST_. Pearson correlations for A) genetic *F*_ST_ and geographic distance, B) cultural *F*_ST_ and geographic distance, C) genetic *F*_ST_ and cultural *F*_ST_, D) genetic *F*_ST_ and linguistic distance, E) cultural *F*_ST_ and linguistic distance, and F) geographic distance and linguistic distance. R corresponds to the correlation coefficient; p corresponds to the p-value.

## Discussion

In this study, we generated genome-wide SNP genotype data and investigated the extent to which geographic and cultural processes shape genetic variation within and among four pastoral populations in northern Kenya. We sampled across multiple layers of social organization - ethnolinguistic groups, clans, and territorial sections - finding that geography, rather than cultural processes, predominantly shape patterns of genetic variation in northern Kenya.

Among ethnolinguistic groups, we observed a clinal pattern of variation with a lack of discrete clustering, particularly between the Rendille and Samburu, and comparatively high levels of intermarriage between them. These results suggest ongoing gene flow between the Rendille and Samburu, the two most closely geographically located groups, but not the most culturally similar. Previous literature has noted intermarriage between the Rendille and Samburu (Spencer, 2012) and relatively higher levels of cooperation (Handley & Mathew, 2020). Genetic clustering of Cushitic and Nilo-Saharan speaking groups (of which the Rendille and Samburu are a part of, respectively) has previously been observed, supporting evidence of gene flow between these larger linguistic groups (Tishkoff et al., 2009). Our findings confirm previous genetic and cultural observations and provide an example in humans where genes are shared between different ethnolinguistic groups at a local geographic scale.

Though we found a clinal pattern of variation, the Waso Borana and Turkana formed fairly discrete clusters of unique genetic variation. Strikingly, in the individuals we sampled, no intermarriage was reported in the Turkana at all, and for Waso Borana, there was no intermarriage with the other three ethnolinguistic groups. These results suggest isolation in Waso Borana and Turkana from the ethnolinguistic groups sampled in this study. However, it is possible our sampling locations may be driving part of these observations. The regions that the Waso Borana inhabit border the other ethnolinguistic groups; however, we sampled individuals from the Merti region, which is an interior region of the Waso Borana territory. Likewise, the Turkana individuals we sampled were from the north and west regions of Turkana that do not directly border the other groups. We speculate that although these are nomadic groups that can traverse large distances, intermixing may occur in boundary regions rather than in interior regions. Future studies including individuals from both interior and border regions may shed light on this.

Perhaps one of the most intriguing results in this study was the observed genetic substructuring within Turkana based on territorial section affiliation. Individuals from the Ngibochoros territorial section are further geographically from the individuals sampled from the other territorial sections, and our results suggest that this geographic separation results in high genetic differentiation between individuals from Ngibochoros with individuals from Kwatela and Ngiyapakuno. Because there is no cultural barrier to Turkana individuals marrying individuals from different territorial sections (i.e., clan level exogamy) and because there is extensive migration in dry season across territorial section boundaries in these nomadic groups, we did not expect to observe genetic substructuring within the Turkana. Rather, we were expecting the Turkana to be largely homogenous, similar to what we observed within the other three ethnolinguistic groups we sampled in this study. The Turkana are however the most populous of the four ethnic groups, numbering approximately 1 million individuals, and having the largest geographic span. It is possible that although there is a shared cultural identity over this larger area, interpersonal interactions and co-mingling between distant Turkana territorial sections are limited. Genetic substructuring has been observed across humans on larger geographic scales (i.e., (Alsmadi et al., 2013; Bryc et al., 2010; Jakkula et al., 2008; Salmela et al., 2011; Tian et al., 2008; Tishkoff et al., 2009; Xu et al., 2009)); however, due to a lack of dense sampling within individual populations across Africa, genetic substructuring within a single ethnolinguistic group has not been widely observed within the continent, nor indeed within as small of a region as we investigate here. Because underlying genetic structure can have implications for case-control studies (Price, Zaitlen, Reich, & Patterson, 2010), our results highlight the importance of accounting for fine-scale substructuring in future genomic studies, particularly with the Turkana, and emphasize the continued importance of characterizing genetic structure across globally diverse human populations.

Within ethnolinguistic groups, we found that Y chromosome haplotypes do not consistently cluster by natal clan affiliation, suggesting that patrilineality may not have a strong impact on patterns of male-specific genetic variation in northern Kenya pastoral populations. Previous research has found that, in groups with patrilineal descent, like pastoralists in Central Asia (Chaix et al., 2007) and the Bimoba in Ghana (Sanchez-Faddeev et al., 2013), males from the same clan have identical or similar Y haplotypes. However, this is not always the case, as seen in tribal Yemen, where Y haplotypes do not clearly cluster by clan (Raaum, Al-Meeri, & Mulligan, 2013). A possible explanation for our finding is that cultural conception of fatherhood, and therefore clan affiliation, does not always correspond with who one’s biological father is. For example, in these groups, offspring from unofficial marriages - unions in which the bride price has not been paid - take on their mother’s clan. This would result in a mismatch in clan assignment for these offspring. Adoption can also result in a mismatch in clan affiliation. Adoption is known to occur in Turkana and an adopted child takes on the clan of their adopted father. Overall, these results highlight that patrilineal descent groups do not always correspond with genetic patriline.

We found that genetic differentiation was highest between ethnolinguistic groups separated by the largest geographic distances, suggesting that geography primarily impacts patterns of genetic differentiation among northern Kenyan populations. Previous studies of genetic structure among human populations in Africa have found correspondence between genetic structure and linguistic affiliation and/or geography, with some studies reporting correspondence of genetic structure predominantly with linguistic affiliations (Bryc et al., 2010; Tishkoff et al., 2009), while others found that patterns of genetic structure predominantly mirror geography and ecological barriers (Babiker, Schlebusch, Hassan, & Jakobsson, 2011; Uren et al., 2016). For northern Kenyan human populations, our results suggest that geography primarily shapes the observed patterns of genetic differentiation.

Though genetic differentiation primarily paralleled geographic distances, for some groups in our study, the pattern of genetic differentiation secondarily paralleled linguistic relationships. Specifically, we found that Turkana and Samburu had lower *F*_ST_ than the Turkana and Rendille, despite the former being geographically more distant from one another than Turkana and Rendille. The close genetic relationship between Turkana and Samburu compared to Turkana and Rendille could be due to shared Nilo-Saharan ancestry between the Turkana and Samburu but could also be the result of sampling from areas not directly bordering each group. Although not commonplace, the Turkana are known to intermarry with both the Rendille and Samburu, and likely occurs in regions bordering each group. Given our sampling strategy, we were unable to assess the extent of gene flow in regions directly bordering each group and if this differs from interior regions. Therefore, though our results suggest that patterns of genetic differentiation may be secondarily influenced by local linguistic affiliations for populations with similar geographic distances, additional sampling across the entire region of each ethnolinguistic group may be needed to validate this finding.

Lastly, we found that cultural differentiation does not predict genetic differentiation among neighboring populations in northern Kenya. Previous studies have used and compared linguistic or cultural distance to genetic distance to understand the extent to which genes and culture/language travel among human populations, with examples in human history where genes and culture/language have been shown to travel together (i.e., (Filippo, de Filippo, Bostoen, Stoneking, & Pakendorf, 2012; Hewlett, De Silvestri, & Guglielmino, 2002; Hunley et al., 2008; Karafet et al., 2016; Lansing et al., 2007)) and others where spoken language has been shown to have no effect on genetic structure (Veeramah et al., 2010). Here, we used cultural *F*_ST_ to test whether genes and culture travel together on a local geographic scale and find that cultural *F*_ST_ and genetic *F*_ST_ are not correlated with each other among northern Kenya pastoralists. We caution against the overinterpretation of this result, however, due to the limited number of groups sampled here. We anticipate this metric of cultural similarity will be of interest for future studies aimed at assessing questions of the movement of genes and culture in humans on both larger and local geographic scales. Taken together with the other results in this study, our findings suggest that geographic proximity, not cultural similarity, may provide a better explanation for the observed patterns of genetic variation among these groups.

## Acknowledgments

This work was funded by the John Templeton Foundation (grant no. 48952) to SM, the National Institute of General Medical Sciences of the National Institutes of Health R35GM124827 to MAW, and The Graduate College at Arizona State University, Achievement Rewards for College Scientists (ARCS) Foundation Phoenix Chapter as a Pierson Scholar, and Arizona State University Chapter Sigma Xi to AMTO. The authors acknowledge Research Computing at Arizona State University for providing high-performance computing resources that have contributed to the research results. The National Museums of Kenya provided institutional support to conduct the research in Kenya. We thank our field research assistants for translating the questionnaires and aiding with data collection: Ekiru Carlystus, Amuria Lotiira, Chegem Muya, Gilbert Topos, Dismas Lomelu, Mohamed Noor Guyo, Abdi Wario, Paul Leramato, Damaris Lekilaui, Julius Longonyek, Sinyati Lesowapir, Rafael Letele, Simon Harugura, Benson Morsa, Ejere Ballo, and Lebo Parkeri. We also thank our participants and host communities for their hospitality and for their continued support in this project.

## Authors Contributions

AMTO, CH, ACS, SM, and MAW conceived the study. AMTO and MAW designed the methodology. SM and CH collected the samples and demographic data. AMTO and EKH extracted DNA and prepared samples for genotyping. SM, CH, and AMTO processed and cleaned the demographic data. AMTO performed all formal analyses and led the writing of the manuscript. All authors contributed critically to writing and editing the drafts and gave final approval for publication.

